# Genotype-Dependent Response to Water Deficit: Increases in Maize Cell Wall Digestibility Occurs Through Reducing Both p-Coumaric Acid and Lignification of the Rind

**DOI:** 10.1101/2025.01.27.635197

**Authors:** Ana López-Malvar, Oscar Main, Sophie Guillaume, Marie-Pierre Jacquemot, Florence Meunier, Pedro Revilla, Rogelio Santiago, Valerie Mechin, Matthieu Reymond

**Affiliations:** Facultad de Biología, Departamento de Biología Vegetal y Ciencias del Suelo, Agrobiología Ambiental, Calidad de Suelos y Plantas, Universidad de Vigo, As Lagoas Marcosende, 36310 Vigo, Spain, Unidad Asociada a la Misión Biológica de Galicia (CSIC).; Université Paris-Saclay, INRAE, AgroParisTech, Institut Jean-Pierre Bourgin for Plant Sciences (IJPB), 78000, Versailles, France.; IATE, University of Montpellier, INRAE, Institut Agro, Montpellier, France.; Unité Expérimentale DiaScope, INRAE, Mauguio, France.; Misión Biológica de Galicia (CSIC), Pontevedra, Spain; UMR AGAP Institute, University of Montpellier, CIRAD, INRAE, Institut Agro, Montpellier, France

**Author notes:** Ana López-Malvar; Oscar Main; Sophie Guillaume; Marie-Pierre Jacquemot; Florence Meunier; Pedro Revilla; Rogelio Santiago; Valerie Mechin; Matthieu Reymond.

**Keywords:** Keywords: drought, maize, p-coumaric acid, lignification, histology, cell wall digestibility

## Abstract

The compositional dynamics of the cell wall are influenced by drought, and it has been demonstrated that water deficit induces significant changes in its main components. Moreover, changes in cell wall concentration and distribution in response to water deficit affect maize degradability. This study presents a histological and biochemical analysis of thirteen maize inbred lines, evaluated over two years in Pobra de Brollón (Spain) and Mauguio (France) under contrasting water availability conditions. Our aim was to investigate the environmental and genotypic impacts on histological and biochemical profiles, to assess in vitro cell wall digestibility under water deficit, and to explore how these responses relate to changes in cell wall composition and structure. Overall, we observed greater concentrations of p-coumaric acid under control conditions, with significant decreases in stressed conditions at each location. Histologically, we found an increase in non-lignified tissues under water deficit conditions across all tissues and in both locations. In terms of in vitro cell wall digestibility (IVCWRD), significant increases were detected in response to water deficit. Additionally, genotype-dependent response patterns were evident, revealing two distinct behavioural groups. Notably, in plastic genotypes, increases in IVCWRD in response to water deficit were concomitant to reductions in p-coumaric acid content and a decrease in red-stained lignified tissues in the rind. This study emphasizes the complex, genotype-dependent responses to water deficit, underscoring the important roles of plasticity and stability in shaping the impact on maize cell wall digestibility; paving the way to breed for adapted genotypes to face climate changes.

**Highlight:** Increased digestibility in response to water deficit is genotype-dependent and linked to reductions in p-coumaric acid content and lower lignification of the rind

## Introduction

In the current climate change context, drought and rising temperatures are two of the biggest challenges for crops. Climatic models predict increases in average temperature, alteration in intensity, severity, and frequency of long-term drought conditions, and a higher risk of heat stress. Maize is one of the most important crops worldwide, and its production, measured as grain yield, is affected by drought at all developmental stages, mainly affecting grain yield and grain quality (Reynolds and Langridge, 2016). Therefore, improving drought tolerance is an important goal of maize breeding.

Not only the grain fraction is affected by drought conditions, but also the compositional dynamics of the cell wall. It has been demonstrated that water deficit induces significant changes in the main components of the cell wall. Emerson *et al.,* (2014) showed a reduction of the cellulose, lignin, and hemicellulose content in maize stover. Similarly increases in β-O-4-linked H lignin subunits in response to water deficit, while lignin and p-coumaric acid contents were reduced have also been reported (El Hage *et al.,* 2018; Virlouvet *et al.,* 2019). Moreover, Álvarez *et al*. (2008) found, under drought conditions, changes in compounds derived from the phenylpropanoid pathway. They argued that accumulating the derived monolignols in the xylem sap may suggest decreases in lignin biosynthesis. They also found increases in peroxidase activity, which may indicate greater cross-linking of cell wall components in response to water deficiency (Passardi *et al.,* 2004).

The quantitative importance of lignin in the cell wall, their structure, and the cross-linkages between cell wall components have variable detrimental effects on cell wall carbohydrate degradation (Barrière *et al.,* 2004; Grabber *et al.,* 2004; Ralph *et al.,* 2004). The changes in cell wall concentration and distribution described in response to water deficit affect cell wall degradability. In water deficit scenarios, an increase in cell wall degradability was associated with reductions in lignin and p-coumaric content within the wall and its preferential localization in cortical regions (El Hage *et al.,* 2018). Furthermore, as water stress increased, agronomic performance declined gradually. However, the digestibility of dry matter (DM) and cell wall (CW) increased consistently regardless of the severity of water stress, accompanied by cell wall composition and structure modifications (El Hage *et al.,* 2018, 2021; Virlouvet *et al.,* 2019; Main *et al.,*2023)

Here, we present a histological and biochemical study of thirteen maize inbred lines evaluated over two years in two locations (Pobra de Brollón, Spain, and Mauguio, France) under contrasting water availability scenarios. Our objectives are to investigate the environmental and genetic impacts on histological and biochemical profiles, assess the response of in vitro cell wall digestibility to water deficit, and explore how this response is linked to changes in cell wall composition structure and distribution.

## Material and Methods

### Vegetal Material and Experimental Design

A set of 13 inbred lines was sown in two locations fields in 2022 and 2023: Centro de Investigaciones Agrarias de Mabegondo-CIAM (Pobra de Brollón, GPS, N: 42’ 33’’15.23, W -7’ 23’’ 18.59) and DiaScope Experimental Unit (Mauguio; GPS, N: 43’36”52.438/E: 3’58”34.419). The 13 inbred lines were selected based on prior experience: inbred lines from Spanish germplasm banks EP1, EPD1, EPD6, EP42, EP105, CO384, PB130 and W64A were chosen due to their differences in cell wall composition (López-Malvar *et al.,* 2021), while inbred lines from French germplasm banks F4, F7, F252, F7019 and F7025 were selected to represent a wide range of lignin content and cell wall digestibility (El Hage *et al.,*2018).

In Pobra de Brollón, the inbred lines were evaluated following a split-plot design. Each irrigation condition (control and drought), comprised three blocks. Each experimental plot consisted of two rows, each row with 20 kernel hills planted manually, spacing between consecutive hills in a row being 0.22 m and 0.75 m between rows, obtaining a final density of ∼60.000 plants. ha^−1^. Local agronomical practices were fulfilled. The two irrigation conditions were implemented by using pairs of tensiometers in each block per condition at - 40cm depth, and pluviometers. A drip irrigation system was applied in the control blocks (PBWW) every 1.5 m, to achieve uniform irrigation. Irrigation was scheduled to water three days a week based on the water availability of the experimental field; resulting in 26 mm/week. In the water deficit blocks (PBWD), no irrigation was applied throughout the development of the experiment.

Similarly, at Mauguio, the inbred lines were evaluated following a randomized block design within each irrigation condition in an incomplete Latin square. Each irrigation condition comprised three blocks. Each experimental plot consisted of two rows per condition, spacing between consecutive hills in a row being 0.19 m, obtaining a final density of ∼95.000 plants. ha^−1^, with all conditions separated by a 20 m buffer area to avoid accidental irrigation of deficit conditions. The three irrigation conditions were implemented by using pairs of tensiometers in each block per condition at -30 and -60 cm depth, and pluviometers. The well-watered condition (MWW) was irrigated with ramp irrigation three times a week with 20 mm of irrigation, a moderate water deficit condition (MWD1) with ramp irrigation of 15 mm when hydric tension reaches –125 kPa at -30cm, a severe water deficit condition (MWD) with ramp irrigation of 13 mm when hydric tension reaches –300 kPa at -30 cm.

To assess the agronomic effects of water stress, plant height was measured at silage stage. Plant height was recorded as the average height (in cm) of five plants per plot. Measurements were taken from the base of the plant to the flag leaf.

### Cell Wall Biochemical analyses

From each plot, at silage stage (the milky to pasty grain transition), four representative plants without the ear were weighed and crushed. A representative sample of fresh stover was weighed and then dried at 55 °C in a stove and again weighed after 72h to estimate the dry matter content. The dry stover samples were ground in a Wiley mill with a 1 mm screen for biochemical analysis.

Cell wall residue was extracted by the Soxhlet water/ethanol method (Effland, 1977). In Vitro Cell Wall Residue Digestibility (IVCWRD) was determined following the methodology described by Lopez-Marnet *et al.,* (2021). The procedure involves an initial pre-treatment step, where 30 mg of dry matter is incubated in a 0.1 N HCl solution containing 2 g/L of pepsin at 40 °C for 24 hours. After this, the sample undergoes a neutralization process, followed by enzymatic hydrolysis using an Onozuka R10 cellulase solution (1 mg/mL) at 40 °C for another 24 hours. Lignin content was determined by treating approximately 7 to 8 mg of extracted cell wall (CW) with acetyl bromide, following the acetyl bromide (ABL) method adapted from Fukushima and Hatfield (Fukushima and Hatfield, 2004). Lignin monomeric composition and structure were analyzed by thioacidolysis, where 13 to 15 mg of extracted cell wall (CW) were treated with boron trifluoride and ethanol at 100 °C for 4 hours in an oil bath. The resulting lignin subunits and β-O-4 linkages were quantified using gas chromatography-mass spectrometry (GC–MS), following the method described by Lapierre *et al*. (1986); hydroxycinnamic acids: p-coumaric acid (PCA), Ferulic acid esterified (FAest) and Ferulic acid etherified (FAeth) were obtained after submitting the cell wall residue through mild and severe alkaline hydrolysis as described in Mechin *et al.,* (2000), following extraction adding and internal standard (p-anisic acid) and finally quantified by high-performance liquid chromatography (HPLC) according to Culhaoglu *et al.,* (2011).

### Histological analyses

At the silage stage, the internode below the main ear from three representative plants was harvested from each plot and stored in 70% ethanol; two internodes per plot were considered for further histological analyses.

A 1 cm-long segment was taken from the upper part of each internode, located 1.5 cm below the node. From each segment, a cross-section of 150 µm was obtained using a GSL1 sled-microtome (Gärtner *et al*., 2014) and preserved in 70% ethanol for further staining in FASGA solution (Fucsina, Alcian blue, Safranina, Glicerina and Aqua). The cross-sections were immersed in a FASGA solution, diluted in distilled water (1:8, v/v), and subjected to 24 hours of agitation. Subsequently, they were rinsed with distilled water for another 24 hours under agitation. Using a slide scanner controlled by the Metafer scanning and imaging platform (MetaSystems GmbH, Altlussheim, Germany), an image of each cross-section was obtained with a resolution of 5.17 µm per pixel (Legland *et al*., 2015; El Hage *et al*., 2021; Lopez-Marnet *et al*., 2022; Main *et al.,* 2025 (under review)). The histological profile of the maize internode cross-section images obtained was quantified by applying the automatic segmentation imaging workflow described by Lopez-Marnet *et al.,* (2022) developed using the ImageJ/Fiji platform. This workflow segments maize internode cross-section images into 40 distinct tissues: two tissues in the epidermis, 19 tissues in the rind, 14 tissues in the pith, and 5 tissues in the bundles. The segmentation is achieved by integrating the Hue, Saturation, and Value properties of each pixel along with the pixel’s location in the FASGA-stained cross-section. Based on enzymatic digestion on cross sections, Lopez-Marnet *et al.,* (2022) also attributed a higher (“b” tissues) or lower (“a” tissues) sensitivity to degradation in pith. To simplify the analysis of the 40 identified tissues, and based on our expertise, we grouped them based on lignification profile: lignified tissues stained red by FASGA and non-lignified tissues stained blue by FASGA. Additionally, we categorized them into three main tissue types: rind, medullary, and bundle tissues. This resulted in the following traits: Rind tissues red (RT_R: DRT2, DRT4, DRT5, DRT7, DRT8, DRT9; LRT2, LRT4, LRT5, LRT7, LRT8, LRT9), Rind tissues blue (RT_B: DRT1, DRT3, DRT6; LRT1, LRT3, LRT6), Medullary tissues red (MT_R: MT1, MT4), Medullary tissues blue (MT_B: MT2, MT5), Bundle tissues red (BT_R: BT1, BT3, BT4), and Bundle tissues blue (BT_B: BT2, BT5). For each individual tissue, the pixel surface area was normalized by the total pixel count of its corresponding main tissue category (rind, medullary, or bundle), allowing the calculation of its percentage relative to the total area within that category. It should be noted that out of the 40 tissues identified, 13 were not assigned due to their limited presence in the sections, which made their identification difficult using the FASGA staining method.

### Statistical analyses

Data was statistically analyzed using R version 4.4.2 (2024-10-31 ucrt) (‘R Core Team 2024) and SAS program (SAS Institute, 2015). The conditions within each location were treated as independent, resulting in five irrigation treatments: MWW, MWD1, MWD, PBWW, and PBWD.

To analyze the biochemical data, a generalized linear mixed model (GLIMMIX) was applied using SAS. The model included condition (location-condition combination), genotype, and their interaction (genotype×condition) as fixed effects, and the year was considered as a random effect. Similarly, for histological data, we used a GLIMMIX model and considered fixed effects genotype, condition (location×condition combination), and their interaction. Random effects for histological data were specified for year bloc(year×condition) genotype×year, and genotype×year×condition to account for variability across years, blocks, and their interactions with genotypes and conditions. For both models, wald tests were used to assess the significance of fixed effects. Least square means (LSMeans) were calculated for condition and genotype, and pairwise comparisons were performed using the diff and lines options to identify significant differences. In parallel for all traits, the best linear unbiased estimator (BLUE) for each genotype×condition was calculated.

We performed a Principal Component Analysis on the BLUEs of histological and biochemical traits separately, considering the interaction genotype×condition. In the Principal Component Analysis analysis, IVCWRD was considered as a spectator variable to explore its relationship with the principal components without influencing their calculation. The Principal Component Analysis was conducted using the first two principal components to reduce dimensionality, using the FactoMineR (Le *et al*., 2008) and factoextra (Kassambara and Mundt, 2020) packages in R. To group the genotypes based on their histological and biochemical profiles, we applied hierarchical grouping and k-means grouping. For hierarchical clustering, we computed the distance matrix using the coordinates from the Principal Component Analysis. For k-means clustering, we assigned genotypes and conditions to groups based on the Principal Component Analysis coordinates, using the stats package. The resulting groups were used to create biplots, including ellipses representing each group, facilitating the visualization of genotype×condition distribution within histological and biochemical contexts, with the ggplot2 (Wickham, 2016) package. Group assignments and visual outputs were generated to identify distinct patterns across genotype×condition interactions.

To study the variation of IVCWRD in response to water deficit, we calculated the relative response at the genotype level for each stress condition (MWD, MWD1, PBWD), using the control at each location (MWW and PBWW) as 100%. This allowed us to explore the different response profiles of the evaluated genotype panel. Based on these relative responses, we then examined the relationship between histological and biochemical traits and IVCWRD by applying a multiple linear regression model. In SAS (SAS Institute, 2015), the stepwise method within the PROC REG procedure was used, with variables having a significance value below 0.15 excluded from the model. IVCWRD was treated as the dependent variable, while histological and biochemical traits that showed a significant condition effect in the variance analysis were used as independent variables. Additionally, Pearson correlations and their corresponding p-values were calculated using the ggcorrplot (Kassambara, 2023) package.

## Results

At the agronomic level, plant height exhibited significant variation across treatments in response to water stress; notably, all treatments differed significantly from one another (Supplementary Figure 1). The tallest plants were observed at MWW, while the shortest corresponded to MWD, representing the severe stress conditions in Mauguio. When comparing treatments in both Mauguio and Pobra de Brollón, differences were significant for plant height between control and stress conditions, with consistently taller plants recorded under control conditions. Plant heights recorded at PBWD treatment were consistently lower than those observed at MWD1.

### Increased water deficit leads to reductions in p-coumaric acid, β-O-4 Linkages, and Syringyl Subunits in Lignin

We observed greater concentrations of p-coumaric acid in both PBWW and MWWthat significantly differ for the stress conditions in each location (Figure 1A). The concentrations of p-coumaric acid found in PBWW do not differ from those found in MWD and MWD1 (Figure 1A). Both the β-O-4 linked-subunit S and β-O-4 yield had lower concentrations in Pobra de Brollón compared to all the treatments in Mauguio, conversely in Mauguio we observed a decrease in both the subunit S and β-O-4 yield affected by increased water deficit (Figure 1 B, C). We observed a significant genotype effect for subunit S, β-O-4 yield, and PCA (pvalue = <.0001). However, we didn’t find significant genotype×condition interaction for any of the studied biochemical traits, suggesting that the level of S subunits and β-O-4 yield was constitutively different between the tested lines whatever the environment. No variation at a condition or genotype level was found for lignin content, FAest or FAeth. The identified significant variations for the H and G β-O-4 linked-lignin subunits are linked to geographical differences: concentrations found in Mauguio are higher than those found in Pobra de Brollon. Nonetheless, we did identify a significant genotype effect.

**Figure 1:**
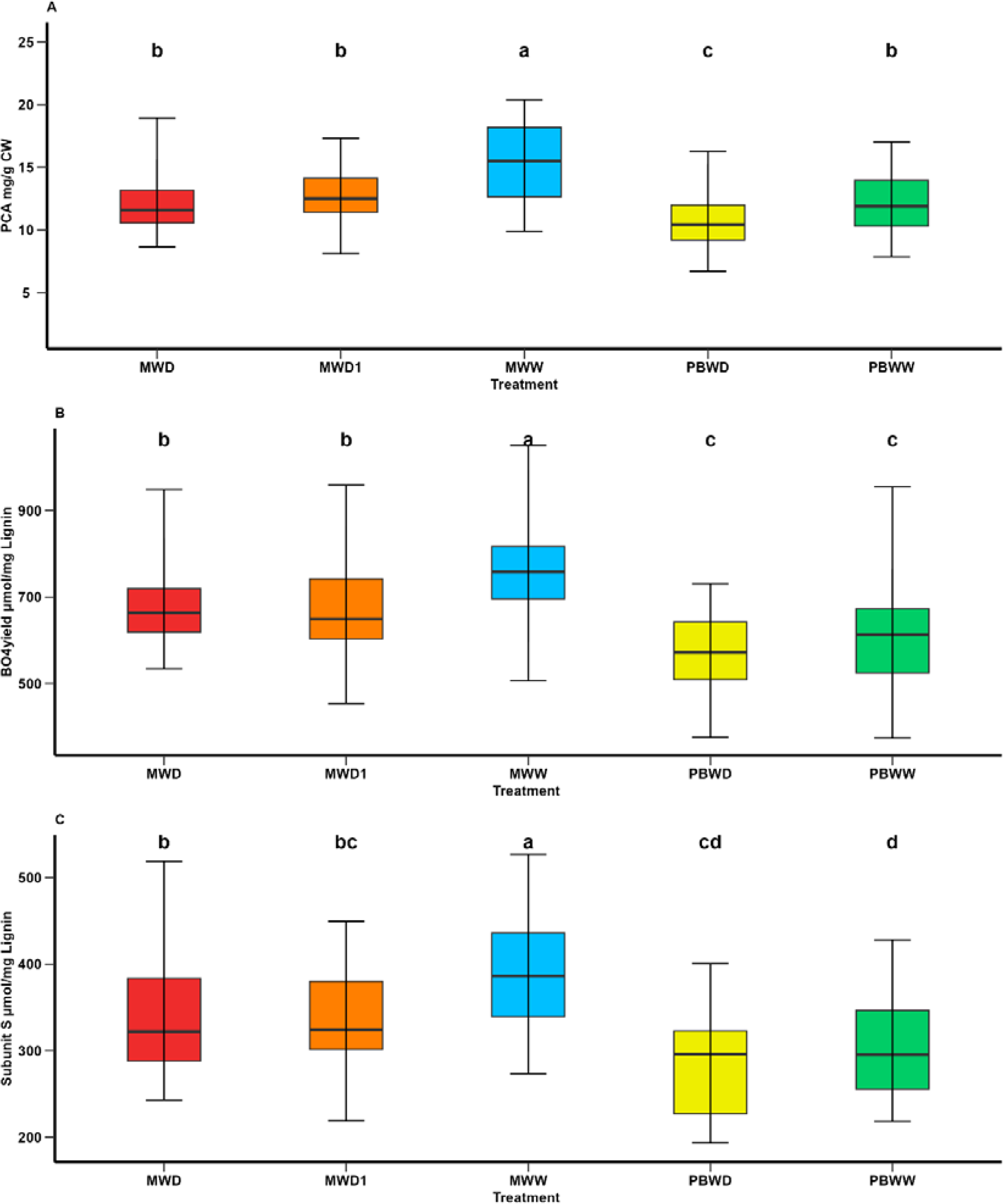
Means comparison of p-coumaric acid and lignin-associated traits under different irrigation treatments. (A) PCA content (mg/g CW); (B) β-O-4-yield (nmol/mg lignin); and (C) Subunits S (nmol/mg lignin). Different letters indicate significant differences between conditions (p < 0.05).

The first two dimensions of the principal component analysis explained 50.3% and 17.1% of the variability, respectively (Figure 2). The first dimension is mainly explained by β-O-4 yield, the subunits S and G, and PCA; while FAest mainly explained the second dimension. When hierarchical and K-means grouping was applied, we obtained 5 groups according to the biochemical patterns observed across genotype×condition interactions, driven by the two dimensions aforementioned, allowing us to differentiate among conditions but also locations. These resulting groups are mostly explained by differences in biochemical profiles. Group 1 (Figure 2, circles) is mainly represented by genotypes in well-water conditions in Mauguio (MWW) which corresponded to samples with greater concentrations of PCA, Subunit S, and β-O-4 yield. Conversely, groups 3 and 4 (Figure 2, squares and crosses) corresponded to genotypes in Pobra de Brollón, in both conditions (PBWW, PBWD), which corresponded to the conditions showing the lowest concentrations of PCA, Subunit S, and β-O-4 yield. Groups 2 and 5 (Figure 2, triangles and square with an X inside) mainly correspond to genotypes in stress conditions in Mauguio (MWD1, MWD), which in general presented lower concentrations of PCA, Subunit S, and β-O-4 yield than MWW (Figure 2, group 1 circles). In this PCA analysis, the variable IVCWRD was treated as a spectator variable, meaning it was projected onto the Principal Component Analysis biplot without contributing to the computation of the principal components. This allowed us to explore its association with the groups and the biochemical traits represented in the Principal Component analysis biplot. In this context, IVCWRD is positioned opposite to those traits that explain the greatest variation along dimension 1 of the Principal Component Analysis (PCA, Subunit S, and β-O-4 yield), highlighting its negative association with the key variables defining this direction of variability. Additionally, IVCWRD is closer to the groups representing genotypes under stress conditions and is only minimally influenced by axis 2, represented by FA, which confirms that this trait has no impact on digestibility variation.

**Figure 2:**
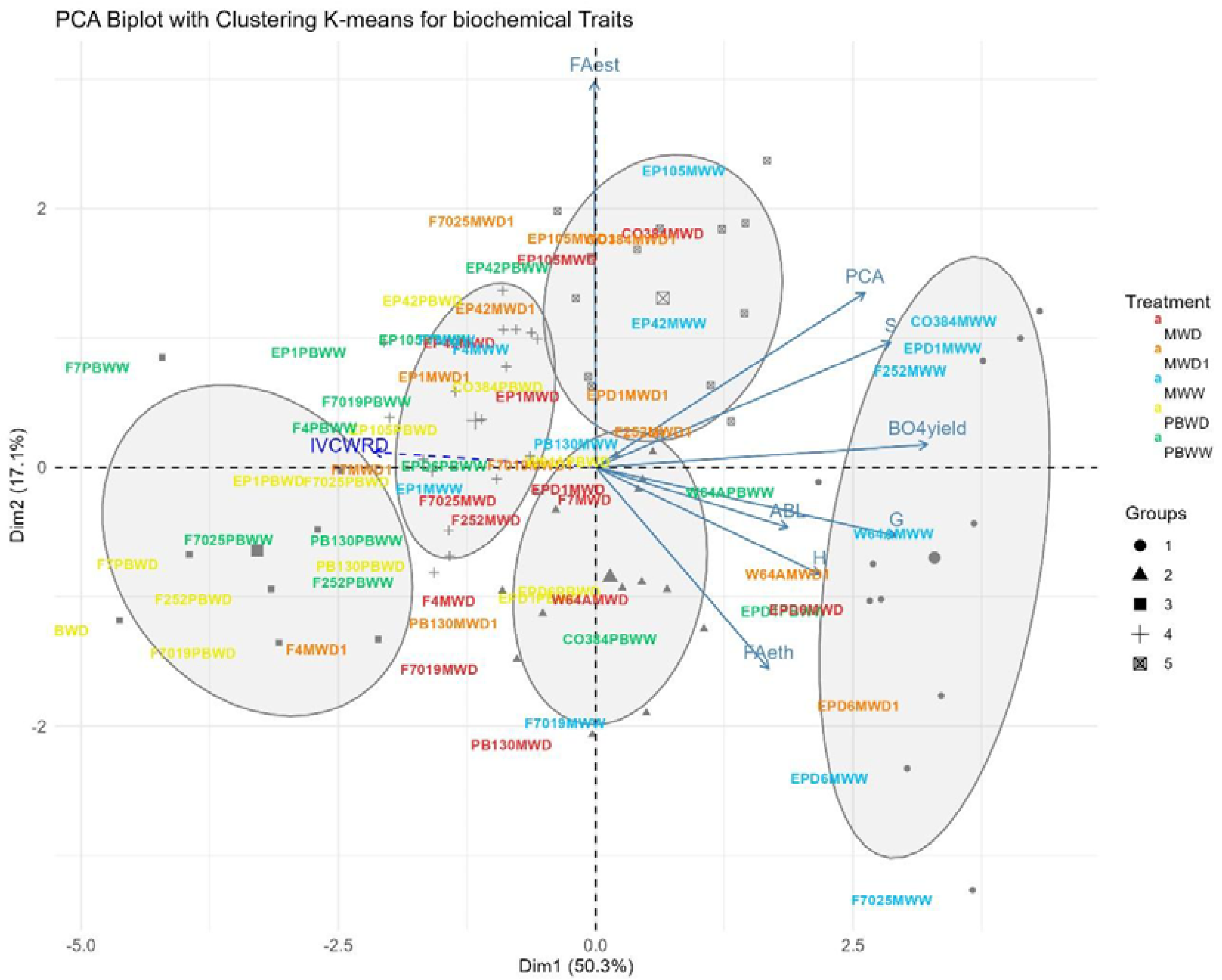
Biplot of the PCA with K-means grouping for biochemical traits. Genotypes in each condition are represented by different colors: MWW in blue, PBWW in green, MWD1 in orange, MWD in red and PBWD in yellow, with groupings further illustrated by ellipses and symbols. Arrows indicate the contributions of variables to the PCA dimensions. IVCWRD, considered as a spectator variable, is represented with a striped arrow in blue.

### Increased non-lignified tissues and genotype-specific patterns as an effect of water deficit on the histological profile

At a histological level, over all the lines, we observed a decrease in lignified tissues in water deficit conditions, at every main tissue level and in both locations (Figure 3). More precisely, for rind tissues, the greatest concentrations of red-stained rind tissues were found in PBWW and MWW, the latest being significantly higher. We found that the red rind values were not significantly different between the stress conditions imposed in Mauguio and Pobra (Figure 3 A). Similarly, lower concentrations of red medullary tissues were found in MWD, differing significantly from the other conditions, and no differences were found among MWD1, MWW, and PBWD (Figure 3 B). We observed significantly lower concentrations of red medullary tissues in PBWD than in PBWW. Regarding the red-stained tissues in the vascular bundles, their concentrations under stress conditions (MWD1, MWD, PBWD) were lower than under well-irrigated conditions (MWW, PBWW), showing significant differences. Significant differences were observed between PBWW and MWW, with the latter exhibiting the greatest concentrations of red-stained bundle tissue (Figure 3 C). For all histological traits, the genotype effect and the genotype×condition interactions were significant.

**Figure 3:**
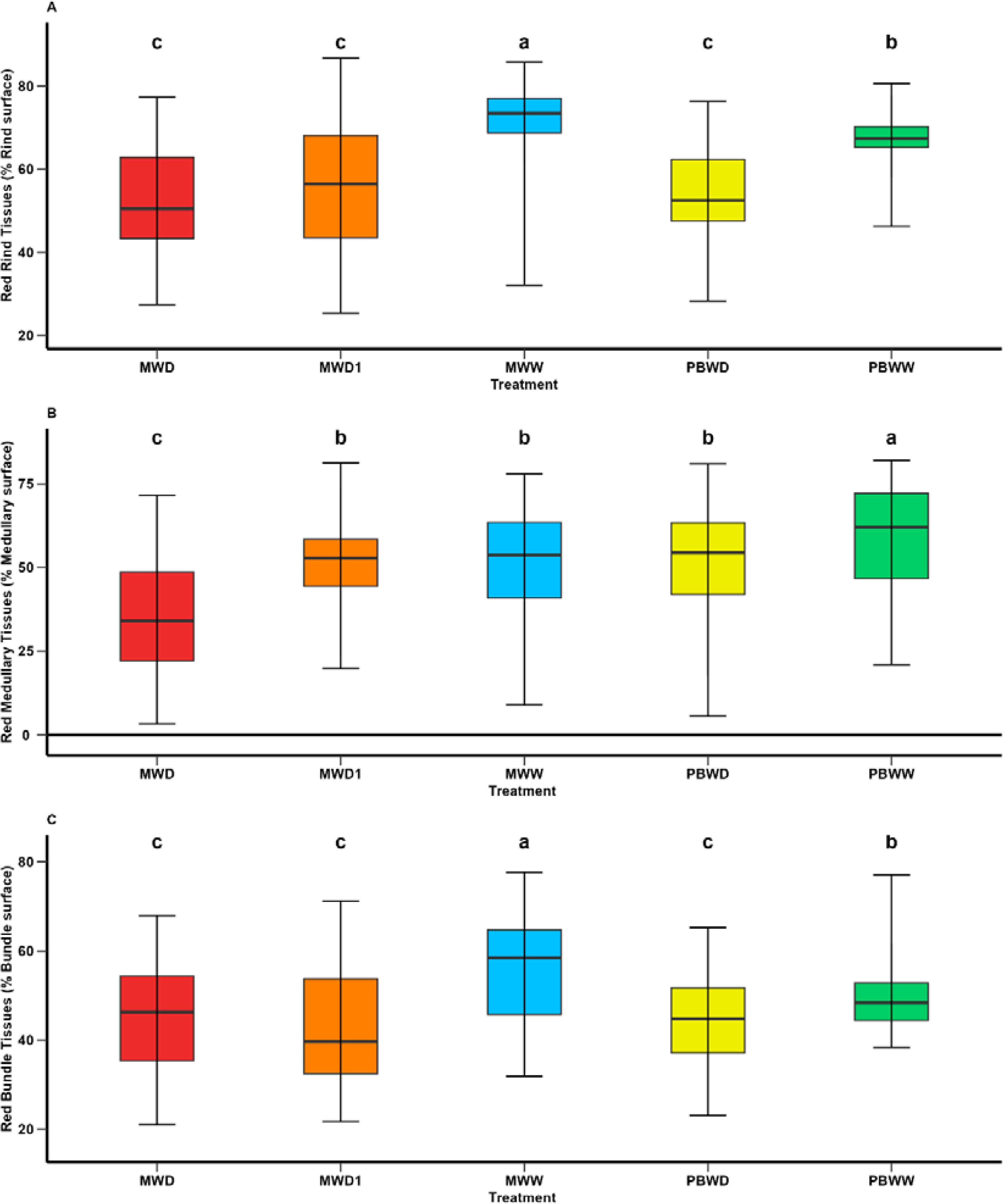
Means comparison of histological traits under different irrigation treatments. (A) Red Rind tissues (% Rind surface); (B) Red Medullary tissues (% Medullary surface); and (C) Red Bundle tissues (% Bundle surface). Different letters indicate significant differences between conditions (p < 0.05).

The first and the second dimensions of the principal component analysis gathering quantified histological traits accounted for 35.1% and 15.9% of the variability, respectively (Figure 4). The first dimension was predominantly influenced by red and blue medullary tissues, while the second dimension was mainly associated with blue rind tissue and red bundle tissues. By applying hierarchical and K-means grouping, three groups were identified based on the histological patterns observed among all inbred lines observed in all environments. The resulting groups highlight the diverse histological profiles present among them. These groups, shaped by the two key dimensions mentioned earlier, effectively distinguish the conditions. In the biplot we differentiated PBWW and MWW grouped in the same group, mainly represented by medullary and rind tissues red and separated from groups 2 and 3 encompassing stress conditions (MWD1, MWD, and PBWD) driven by medullary, rind, and bundle tissues blue (Figure 4). Furthermore, in the biplot, we should highlight particular histological profiles for some of the genotypes. For instance, F7 no matter the conditions always showed a red-lignified profile and all the genotype×condition is grouped in group 3. Conversely, F4 and EP42 in all conditions are grouped in group 1, as no matter the condition these genotypes show a blue-non-lignified profile. In this case, as a spectator variable, IVCWRD is positioned parallel to the blue tissues of both the rind and the blue bundles, with genotypes under stress being positioned closer to it. These traits defined group 1, which included many of the genotypes under severe stress conditions in Mauguio.

**Figure 4:**
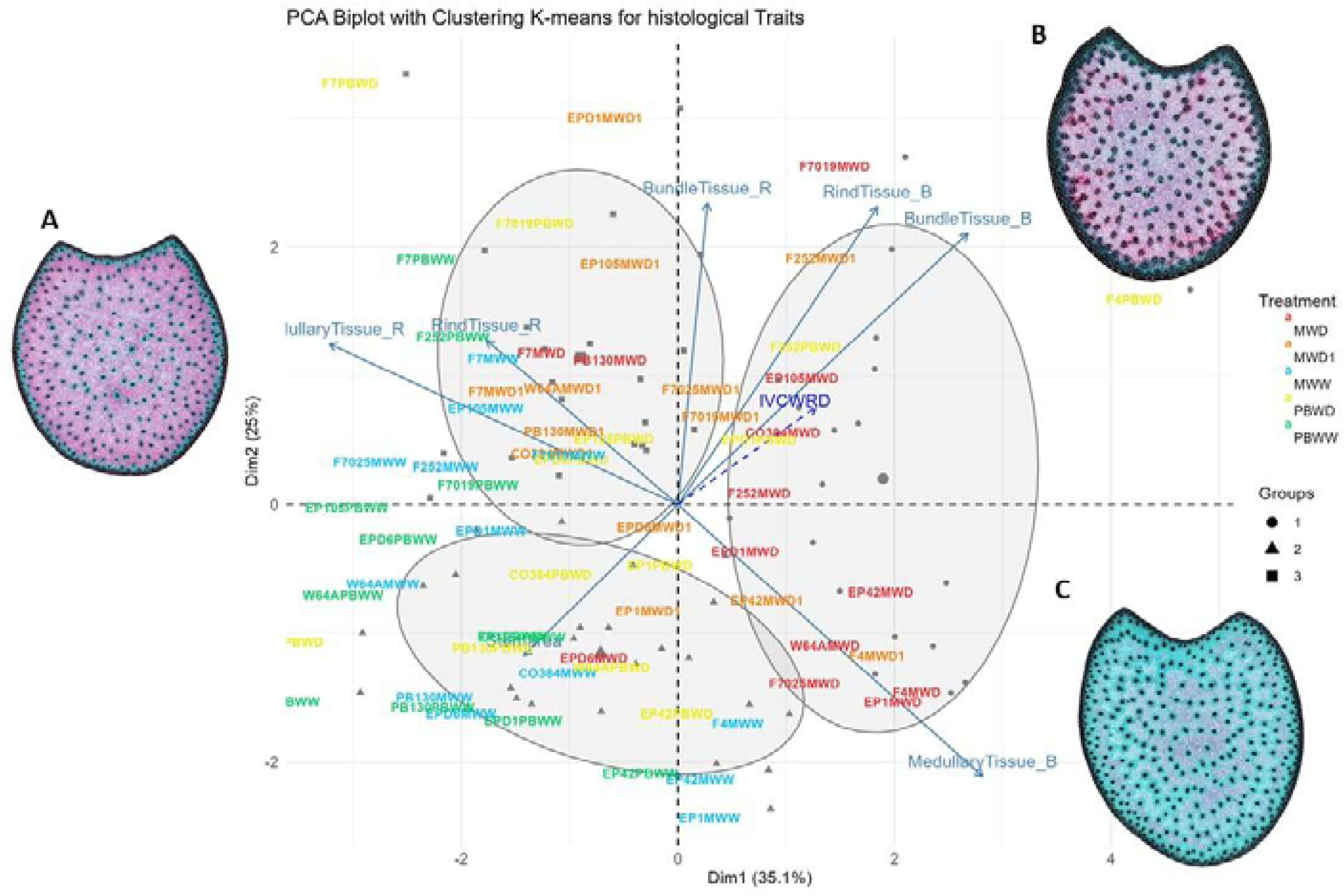
Biplot of the PCA with K-means grouping for histological traits. Genotypes in each condition are represented by different colors: MWW in blue, PBWW in green, MWD1 in orange, MWD in red and PBWD in yellow, with groupings further illustrated by ellipses and symbols. Arrows indicate the contributions of variables to the PCA dimensions. IVCWRD, considered a spectator variable, is represented with a striped arrow in blue. General histological profile in (A) Well Watered conditions, (B) Intermediate Water Deficit, (C) Severe Water Deficit.

### Increased digestibility in response to water deficit is genotype-dependent and linked to reductions in p-coumaric acid content and lower lignification of the rind

In Mauguio, we observed a significant increase in IVCWRD under water deficit conditions, with no differences detected between MWD1 and MWD, in all inbred lines. Conversely, at Pobra de Brollón, no significant variation was observed in response to water stress, as the values for PBWW and PBWD were not significantly different (Figure 5), in all genotypes. Regarding the relative response of each genotype to the different conditions, we can identify different response profiles of IVCWRD behavior in response to water deficit: those genotypes that respond increasing their IVCWRD and those that do not respond (Figure 6). Only the response of the lines in Mauguio was considered for the classification of the profiles since in Pobra de Brollón we did not find significant differences for IVCWRD. As indicated in Figure 6, lines assigned to the red group showed an increase in digestibility between 10-27% relative to the control conditions in Mauguio. Similarly, in Pobra de Brollón, although no differences in digestibility were found between treatments, there was a trend towards an increased response under water deficit, similar to the pattern observed in Mauguio. Specifically, the red group included the following lines: CO384, EP104, EPD1, F252, F7025, PB130 and W64A. In contrast to the red group, the black group includes a set of lines that show more diverse responses under the same conditions. While genotypes in this group achieve a modest increase in digestibility of over 5% (EPD6 and F4), others exhibit variable (F7) or even decreasing trends (EP1), as depicted in Figure 6. Also included in the black group are EP42 and F7019 which show a combination of moderate increases and fluctuating trends, reflecting a unique behavior under the same conditions. Overall, we identified two main behavioral patterns: response and stability. The red group exemplifies the response pattern, characterized by significant increases in digestibility in response to water deficit. In contrast, the black group reflects varying degrees of stability, maintaining lower or negative changes or showing moderate increases combined with unique fluctuating trends. Although no overall significant variation was found for IVCWRD’s response to water deficit in Pobra, we identified some genotype-dependent responses similar to those found in Mauguio.

**Figure 5:**
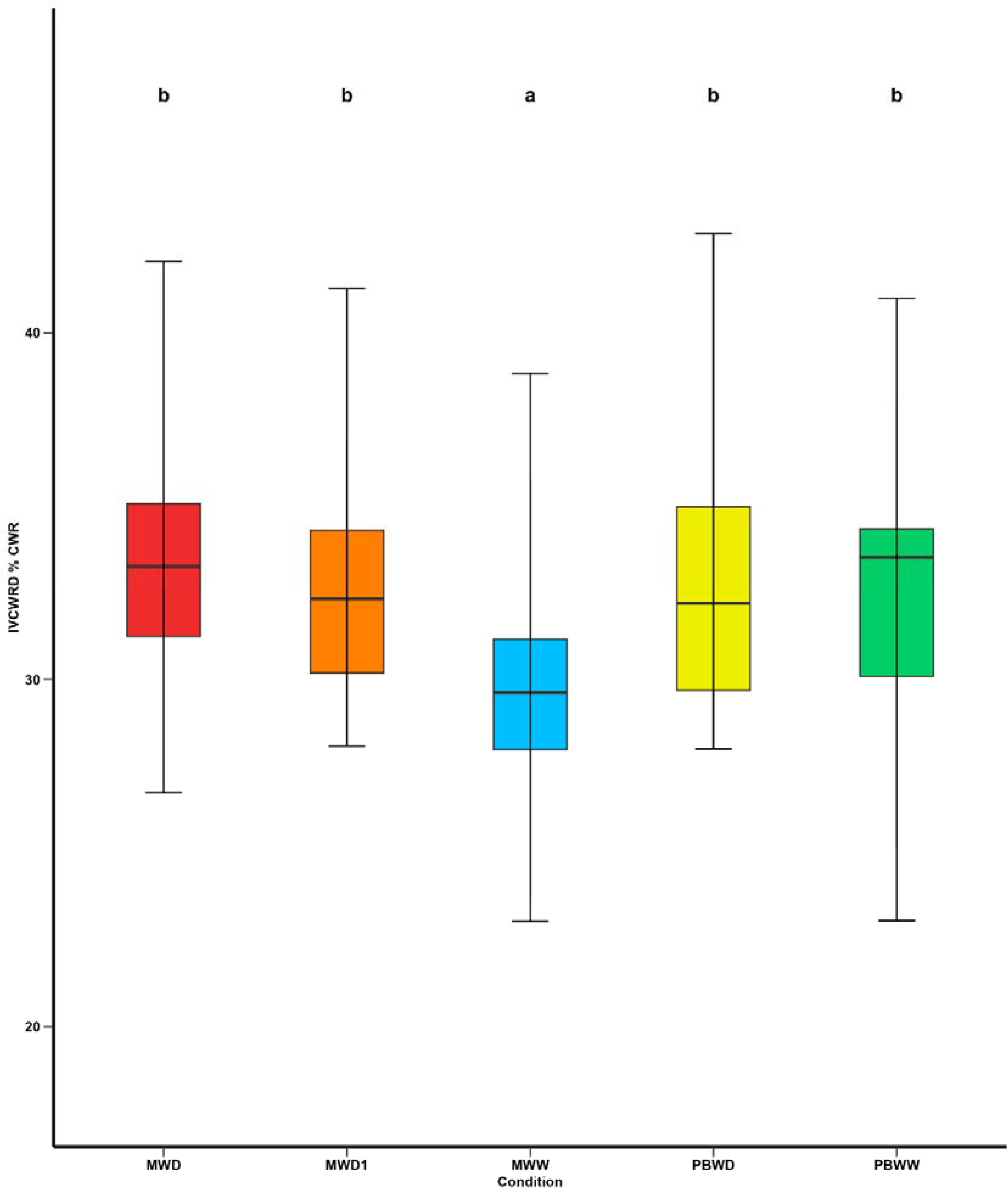
Means comparison of In Vitro Cell Wall Residue Digestibility under different irrigation treatments. Different letters indicate significant differences between conditions (p < 0.05).

**Figure 6:**
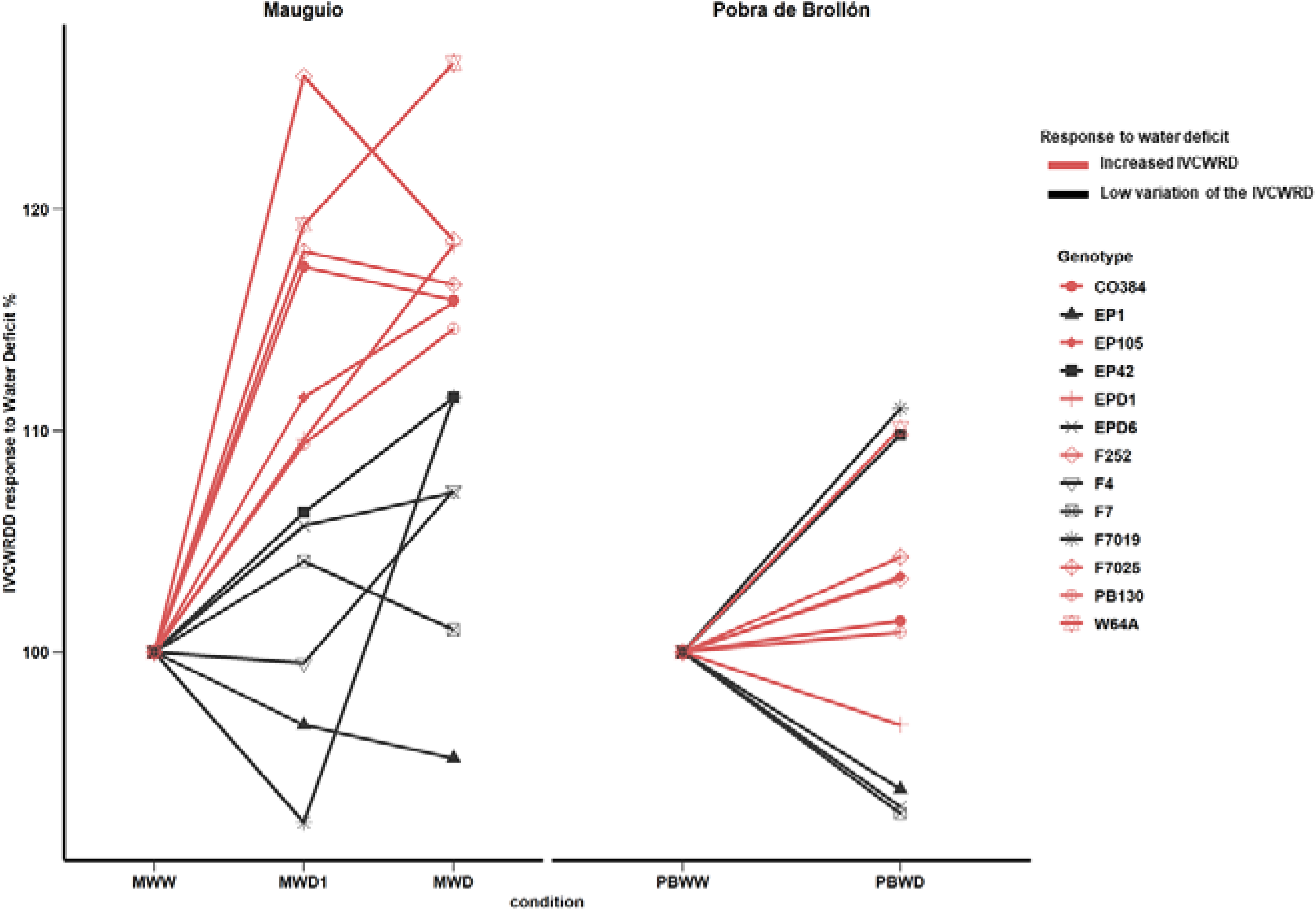
Evolution of In Vitro Cell Wall Residue Digestibility in response to water deficit by genotype; at Mauguio on the right and Pobra de Brollón on the left. Control conditions in each location were considered 100%.

To study if the variations of IVCWRD response are accompanied by variations at a histological and biochemical level, we performed a multiple linear regression model. We found that increases in IVCWRD in response to water deficit are accompanied by decreases in PCA content, explaining 46% of the variability of the response, and with decreases in RT_R, explaining an additional 4% of the variation for IVCWRD response (Table 1).

**Table 1:** Multiple linear regression stepwise model selection and equation for In Vitro Cell Wall Digestibility response to water deficit.

|  |  |  |  |
| --- | --- | --- | --- |
| Model | IVCWRD=148.98-0.36*PCA -0.13*RT_R |  |  |
| Trait | Partial R<sup>2</sup> | Model R<sup>2</sup> | Pr > F |
| PCA | 0.4578 | 0.4578 | <.0001 |
| RT_R | 0.0359 | 0.4936 | 0.1243 |
| IVCWRD: In Vitro Cell Wall Digestibility; PCA: p-coumaric acid; RT_R: Rind Tissues red |  |  |  |

To delve into the relation between the response of both histological and biochemical traits and IVCWRD to water deficit, as well as the genotypic response profile, we explored the relation of the traits under stress conditions (Figure 7). In general, and in agreement with the results obtained in both multiple linear regression and correlations, increases in IVCWRD are linked to reductions in red tissues in the rind and PCA content.

**Figure 7:**
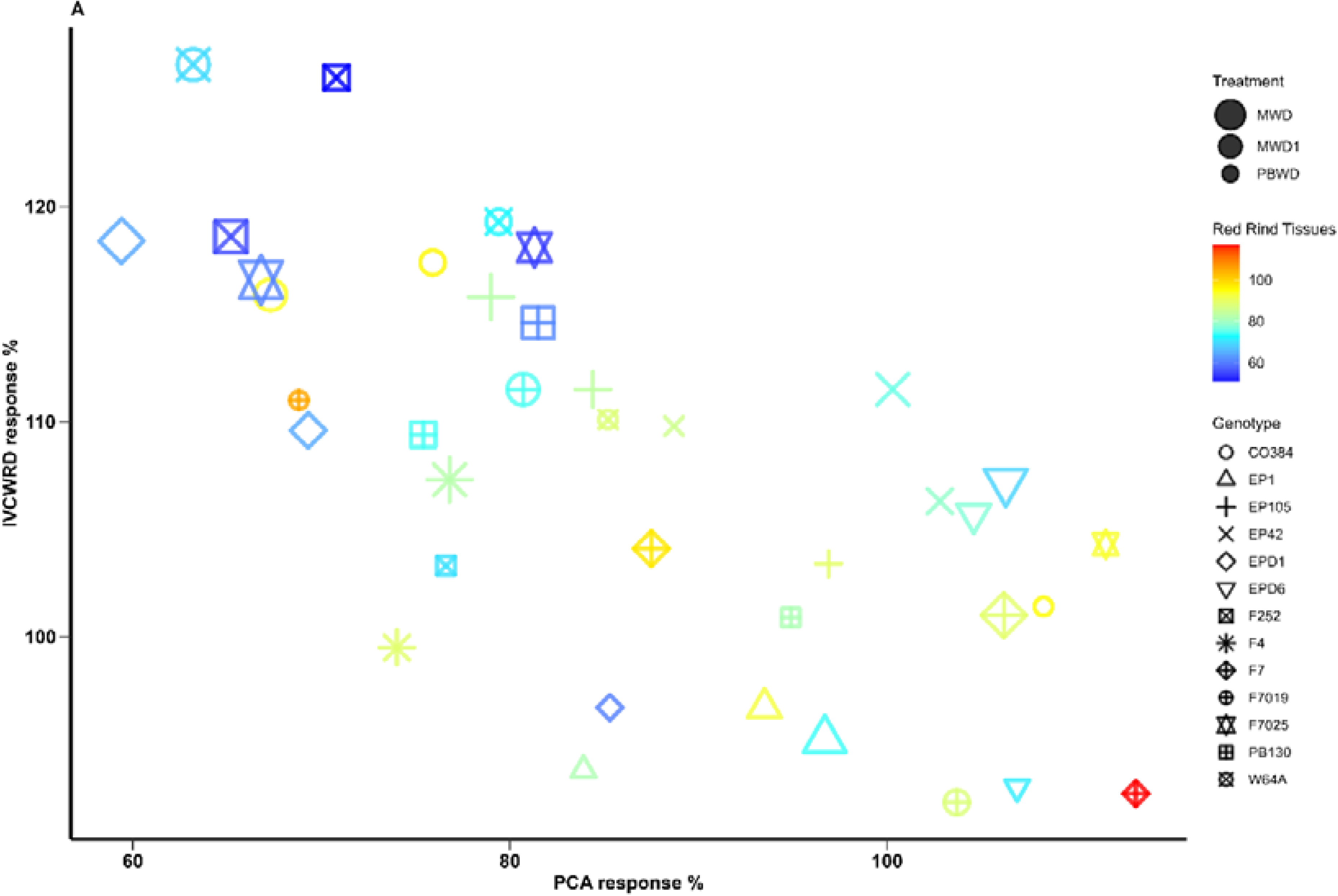
Co-variation between In Vitro Cell Wall Residue Digestibility, PCA and Red Rind tissues in response to water deficit. Red Rind tissues response is represented with a color range and genotypes are represented with different symbols. The treatments are represented in different sizes: MWD > MWD1 > PBWD.

### Water Deficit Induces Shifts in Tissue Composition and Biochemical Traits Linked to Increased Digestibility

We studied the correlation of histological and biochemical traits in response to water deficit, as well as their relationship with IVCWRD response (Figure 8). There were significant patterns of association that highlight the interplay between tissue composition and biochemical properties in response to water deficit (Figure 8). To avoid redundancy, only red-stained tissue responses are presented.

**Figure 8:**
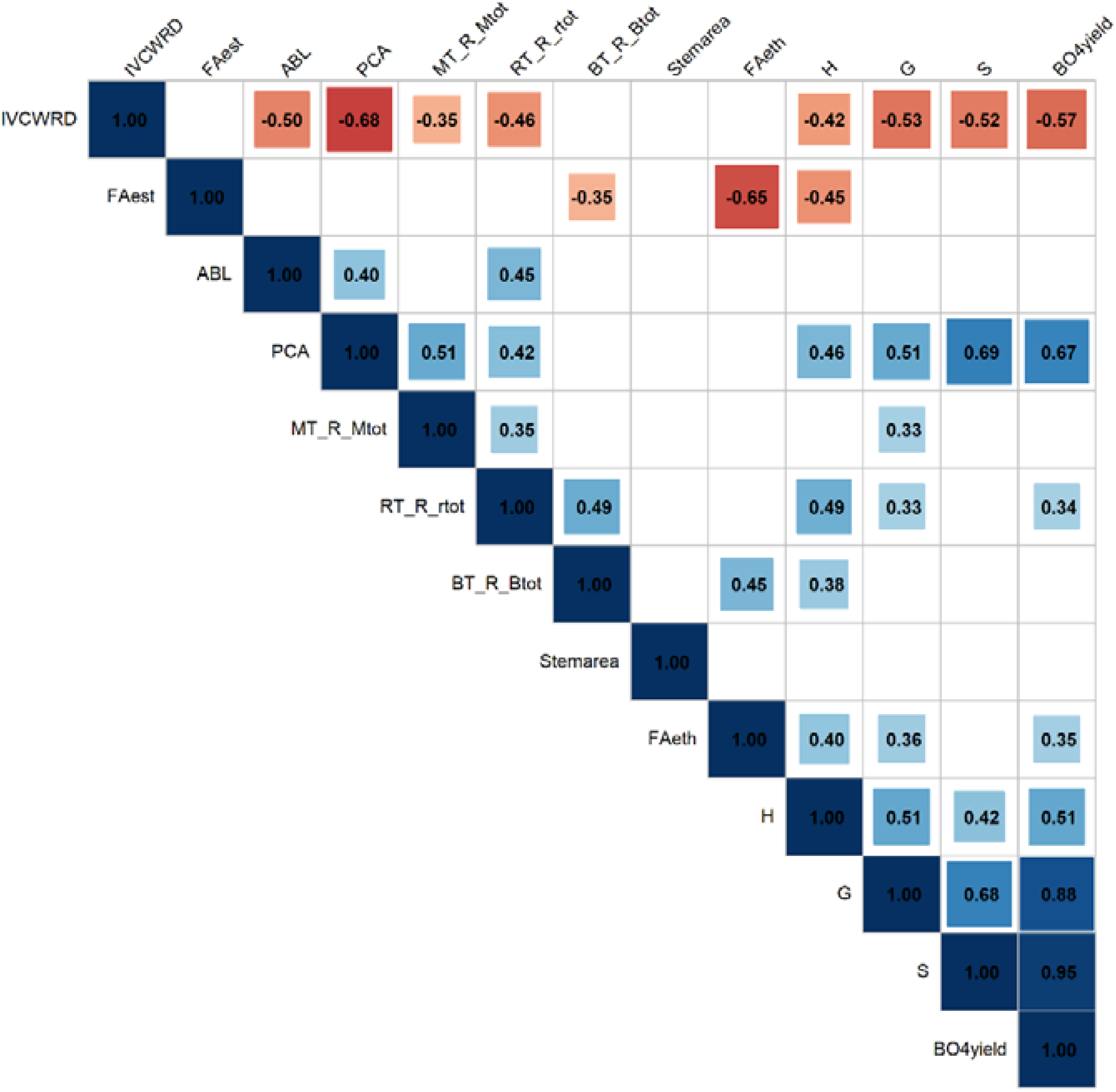
Pearson correlations among biochemical and histological traits in response to water deficit. Only the response in water deficit conditions (MWD, MWD1, PBWD) was considered for the correlations. FAest: Ferulic acid esterified (mg/g CW); FAeth: Ferulic acid etherified (mg/g CW); PCA: p-coumaric acid (mg/g CW); ABL: Acetyl bromide lignin (mg/g CW); H: Subunit H (nmol/mg lignin); S: Subunit S (nmol/mg lignin); G: Subunit G (nmol/mg lignin); βO4yield: β-O-4-yield (nmol/mg lignin); MT_R_Mtot: Red Medullary tissues; RT_R_rtot: Red Rind tissues; BT_R_Btot: Red Bundle tissues; Stemarea: Stem crosssection area.

**Figure 9:**
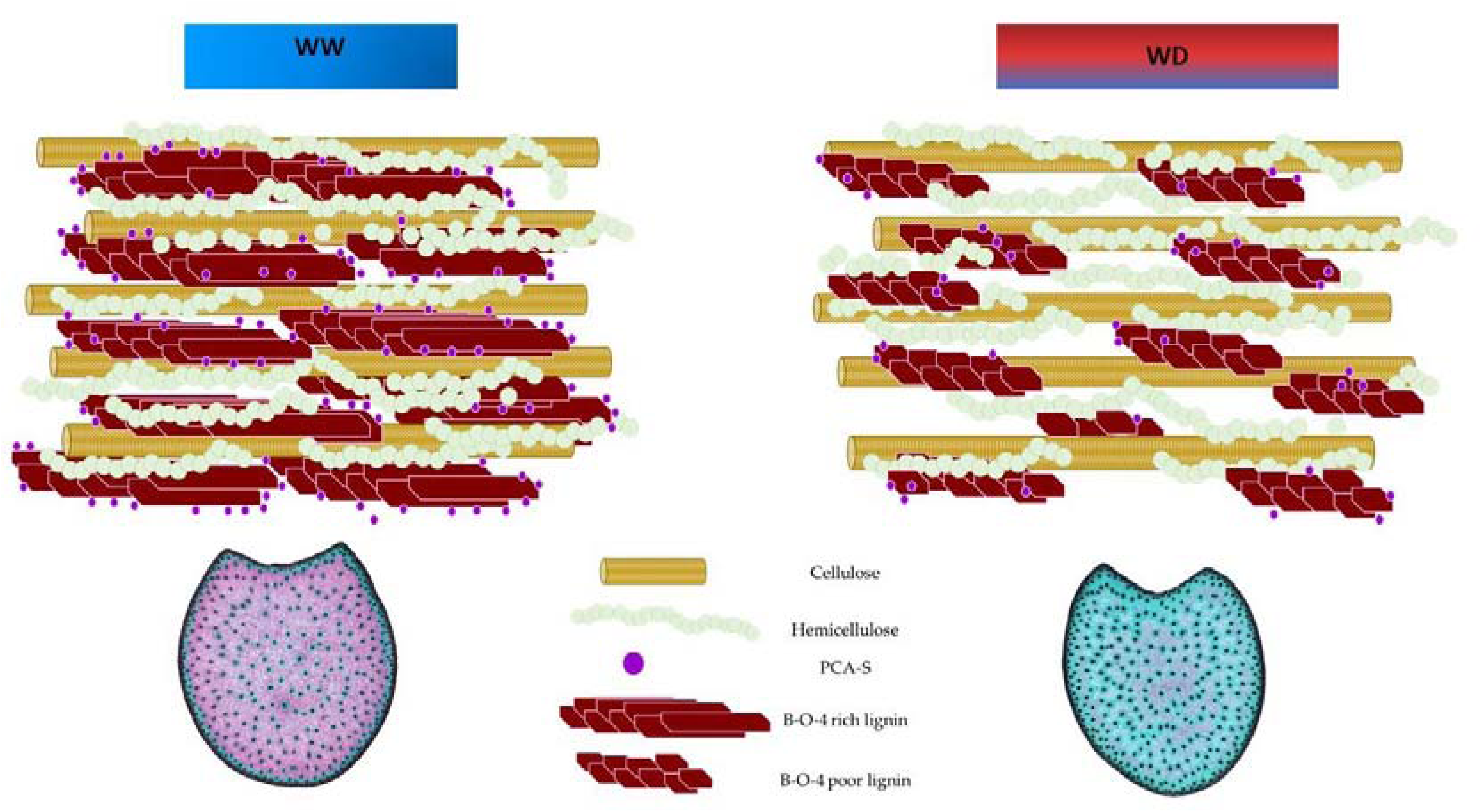
General biochemical and Histological profile of responding genotypes under well-watered (WW) and water deficit (WD) conditions.

Among the histological variables, we found strong and significant correlations between the responses of main tissue types: Red rind tissues response is positively correlated with the responses in the rind and the pith, however, responses of the red tissues of the pith are not correlated with those of the vascular bundles. Notably, there are strong negative correlations between responses of lignified tissues (stained red by FASGA) and non-lignified tissues (stained blue, data not shown). For biochemical variables, we observed significant positive correlations, particularly among responses of lignin-related traits such as PCA, β-O-4 linked S units, and β-O-4 yield, as well as between the lignin subunits themselves and β-O-4 yield. Interestingly, lignin content (ABL) response was only positively correlated with PCA content and Red Rind tissues. Red-stained tissues in the rind and the pith, both showed positive correlations with PCA content and β-O-4 linked subunit G response. Besides, red rind tissues were positively correlated with the response of lignin, subunits H, and β-O-4 yield. Red bundle tissues were positively correlated with subunit H and FAeth. On the other hand, the esterified ferulic acid response was negatively associated with red stained-lignified tissues in the bundles and with FAeth.

For IVCWRD’s response to water deficit, we found strong negative correlations with all the biochemical traits and with the red lignified tissues in the rind and the pith. Particularly noteworthy were the strong negative correlations with PCA, β-O-4 yield, and the S subunit, as well as with red-stained tissues in the rind.

## Discussion

Our study provides a comprehensive analysis of the effects of water deficit on histological and biochemical traits in maize, alongside its impact on in vitro cell wall digestibility (IVCWRD). At the agronomic level, all plants exhibited a clear response to water stress, with plant height decreasing progressively as the severity of stress increased, MWD being the treatment that showed the lowest plant height (Supplementary Figure 1). This is in accordance with other authors who also found decreases in plant height (Emerson *et al*., 2014a; Perrier *et al*., 2017; Main *et al*., 2023). These results confirm the successful implementation of a controlled experimental setup that generated distinct stress levels, effectively mimicking real-world water deficit scenarios and their impact on maize growth. However, the biochemical and histological profiles demonstrated genotype-dependent responses to water stress as well as more pronounced differences due to environmental effects. Similarly, digestibility also varied significantly depending on the genotype and the environment, highlighting the differential adaptability of maize lines to water stress at both the structural and functional levels. These findings underscore the complex interplay between biochemical and histological traits in shaping IVCWRD response under water deficit, offering insights into genotype-specific adaptation strategies. Differences between environments (Mauguio and Pobra de Brollón) are more important for well-watered than for drought conditions; probably due to the more favorable environmental conditions for maize growth in Mauguio.

Histologically, water deficit resulted in reduced lignified tissue across all tissue types, highlighting a shift in tissue composition under stress. The generalized nature of this response was evident across both locations, as demonstrated not only by the variance analysis but also reinforced by the principal components and grouping analysis. Moreover, this response was consistent across the entire histological profile. El Hage *et al.,* (2018), hypothesized differential patterns of lignin deposition as an effect of water deficit, showing a preferential deposition of lignified tissues in the rind in water deficit scenarios. Similarly, the correlations between biochemical and histological responses indicate that, under water-deficit conditions, these reductions in lignin at the rind and medullary level are accompanied by decreases in esterified PCA.

These variations in our dataset in the biochemical profile under water deficit were particularly evident in Mauguio, where levels of PCA, β-O-4 yield and subunits S were higher. In Pobra de Brollón, although baseline concentrations were lower, PCA exhibited an even greater decrease under stress. This pattern aligns with previous studies that highlight the strong interconnection between these traits in their response to water deficit, explaining their concurrent variation (Emerson *et al*., 2014b; El Hage *et al*., 2018, 2021; Virlouvet *et al*., 2019). However, unlike other studies that report changes in total lignin content (El Hage *et al*., 2018, 2021), our results did not show significant differences in lignin levels between water conditions or locations.

In terms of IVCWRD response, significant increases were detected in response to water deficit. Beyond these observations, genotype-dependent response patterns were evident, revealing two distinct behavioural groups. One group exhibited a high level of responsiveness, while the other demonstrated more stable or even declining trends in response to the treatment. Crucially, in plastic genotypes, increases in IVCWRD in response to water deficit, were concomitant with reductions in PCA content and red-stained lignified tissues. Notably, PCA content and the presence of red-stained tissues in the rind together explain 49% of the variability in IVCWRD, with red-stained tissues alone accounting for an additional 4% once PCA content is included in the model. However, the relatively low additional contribution of red-stained tissues (only 4%) can be attributed to their positive correlation with PCA content. These results suggest that PCA content reduction in response to water deficit occurs mainly in the rind. This correlation indicated that changes in PCA content in response to drought, drive the observed variation in red-stained tissues, highlighting its key role in digestibility. Interestingly, the histological and biochemical profiles were tightly interconnected. In this framework, integrating biochemical assessments of dry matter with histological profiling of the stem allows for identifying the specific areas where biochemical changes could occur in response to stress.

Besides, those genotypes showing the greater decrease in PCA are also those showing the strongest response in terms of increased IVCWRD. This is often accompanied by a reduction in red-stained tissues in the rind in these highly plastic genotypes. However, some genotypes that do not respond with increasing IVCWRD significantly reduce their lignification in the rind without showing any increase in digestibility. Specifically, increases of 15-25% IVCWRD in response to water deficit, are associated with decreases of 10-40% in PCA content. However, in the case of red tissues in the rind, a decrease in the lignified tissue surface alone does not guarantee a response increasing IVCWRD unless accompanied by a corresponding reduction in PCA content. That’s the case of the inbred line EPD6 which showed a 20-30% reduction on lignified tissue surface in the rind in response to water deficit without increasing its IVCWRD. This may be explained by the fact that its PCA content does not change significantly in response to stress. The same is true for EP1 and EP42. In contrast, inbred lines EP105 and CO384, which showed plastic behaviour, demonstrated increases in IVCWRD under water stress by reducing PCA content by 15–20%, while showing only minor changes in the lignified tissue surface of the rind. On the other hand, the inbred line F4 reduced its PCA content and lignification in the rind under water-deficit conditions but did not show a significant increase in cell wall digestibility compared to the control. Notably, F4 consistently exhibited the highest digestibility across all water scenarios (data not shown), which may explain its limited response to water stress. Its inherently high digestibility likely leaves less room for improvement/response. In general, variations on those traits in Pobra de Brollón are less evident than for Mauguio, which may explain the lack of significant differences in IVCWRD response to water deficit. The inbred line F4 stood out as a genotype with exceptionally high cell wall digestibility, comparable to that of several brown-midrib (bm3) mutant lines, as noted by Barrière *et al.,* (2017), which inherently presents low lignin and p-coumarate levels, compared to other genotypes. Besides, F4 presents a low lignified parenchyma and vascular bundles. Despite its high digestibility, attempts to introduce bm mutations into its genome did not significantly enhance this trait. This mutation typically impairs the biosynthesis of syringyl units, reducing the S/G ratio to lower values. However, in the case of F4bm3, the S/G ratio remained closer to that of normal lines, with only a slight reduction in syringyl units compared to other bm3 lines. Similarly, in this study, F4’s digestibility remained stable under environmental conditions, such as water-deficit stress, which could increase cell wall digestibility in other genotypes that do respond (Barrière *et al*., 2017).

Most p-coumarate accretion occurs in tandem with lignification. In maize, lignins are acylated (primarily syringyl units) at the γ-position by p-coumarates (Ralph *et al*., 1994). Acylation has a marked influence on the bonding mode of S lignin units, on the spatial organization of lignins, and consequently on their capacity to interact with polysaccharides. It is known that syringyl type lignin form a more linear structure (Kishimoto *et al*., 2005), with little or no branching and with a lesser degree of polymerization that extends further into the secondary wall, protecting a larger proportion of the polysaccharides in the wall from digestion; thus, reducing cell wall digestibility. Building upon this point, Zhang *et al.,* (2019) combined biochemical and histological approaches to characterize cell wall deposition and lignification during maize stem development. Their findings highlighted the spatial and temporal variation in cell wall component deposition and structure across the three main stages of cell wall development: fast deposition of the primary cell walls in whole cross sections; fast deposition of the secondary cell walls; slow deposition of the secondary cell walls in the cortical region. During the secondary cell wall development, rapid deposition of lignin with both C-C and β-O-4 bonds contributed to the fast lignification process. Subsequently, lignin with β-O-4 bonds became predominant during the late stages of lignification, primarily in the rind region. In our study, water deficit conditions were imposed in Mauguio 25 days after sowing. Conversely, in Pobra de Brollón, the absence of irrigation naturally imposed water deficit conditions, particularly as sowing occurred around May 15th, coinciding with the onset of summer and higher temperatures. The strong correlations observed between rind lignification and PCA content response under water deficit suggest that these traits may vary in tandem. At the time the stress was imposed, the pith was fully lignified, as suggested by Zhang *et al*., (2019). However, in the rind, the lignification and deposition of secondary cell walls were still ongoing. As shown in the diagram presented in Figure 10, for those plastic genotypes that responded to drought, under water deficit, we observed a reduction in rind lignification, which was accompanied by decreases in PCA content, S-lignin subunits, and β-O-4 linkage yields. This suggests that the plant may adapt by limiting secondary wall deposition in this region under stress. Such a response highlights the complex biochemical and structural adjustments in maize stems and points to a distinct stress adaptation strategy in these tissues (Figure 10, modified from Amin *et al*., 2014).

**Figure 10:** General biochemical and Histological profile of responding genotypes under well-watered (WW) and water deficit (WD) conditions.

The absence of significant differences in IVCWRD in response to water deficit in Pobra de Brollón, despite reductions in PCA content and decreased lignification in the rind and the pith, can be attributed to the baseline PCA concentrations of the lines in the study in Pobra de Brollón. The PCA content in PBWW did not differ significantly from that in MWD and MWD1, yet a greater reduction in PBWD was still observed. Regarding lignification in the rind, changes in lignin content alone, if not accompanied by corresponding reductions in PCA, were insufficient to enhance digestibility, as previously suggested. Furthermore, in the Pobra de Brollón trial, the levels of syringyl (S) units and the β-O-4 yield were significantly lower than in Mauguio, and these traits were strongly positively correlated with PCA content. These lower concentrations may indicate the presence of a more globular lignin type, as opposed to the linear structures often associated with Syringyl-type lignin. Globular lignin could limit their extension into the secondary wall, reducing the shielding effect on polysaccharides and thereby enhancing cellulose accessibility to enzymes. This structural difference could lead to increased cell wall digestibility (Kishimoto *et al*., 2005). These differences in biochemical composition also result in the genotypes from Pobra de Brollón grouping separately from those in Mauguio in the Principal Components Analysis-based clustering.

Our study highlighted the complex, genotype-dependent responses to water deficit, emphasizing the significant role of plasticity and stability in shaping the impact on maize cell wall digestibility. The observed variation in lignification patterns at the histological level and biochemical traits, particularly the interaction between lignified tissues, PCA levels, and digestibility, provide valuable insights into potential breeding strategies. The presence of distinct behavioral groups—ranging from highly plastic to stable genotypes—suggest different ideotypes depending on water availability conditions. In the absence of water deficit risk, selecting maize varieties that are easily digestible, regardless of their response, would be advisable. However, in high drought-risk conditions, it may be beneficial to select genotypes that exhibit a strong stress response, capitalizing on increased digestibility and all associated traits. Such an approach could pave the way for implementing targeted breeding programs aimed at developing crop varieties better adapted to specific stresses or agroecological contexts. Incorporating this knowledge into breeding programs will provide an effective strategy for developing maize varieties that are not only more resilient to drought but also better suited for sustainable agricultural practices in the face of climate change.

## Acknowledgements

We thank all members of the ’Biomass Quality and Interactions with Drought’ team at IJPB and the ’Maize Genetics and Breeding’ team at Misión Biológica de Galicia for their valuable support throughout the development of this study. We also sincerely acknowledge the DiaScope EU team and the field trial managers from the Pobra de Brollón teams for their dedicated work on the different field trials. The IJPB benefits from the support of Saclay Plant Sciences-SPS (ANR-17-EUR-0007). This work has benefited from the support of IJPB’s imaging and microscopy platform (PO-Cyto).

## Author contributions

MR, VM, PR, RS, and ALM conceived the study. MR, VM, PR, RS, FM participated in the experimental design and carried out the field trials; MR, VM, PR, RS, FM, OM, SG, MPJ and ALM participated in the sample collection; ALM, SG, and MPJ carried out the biochemical determinations, MR, VM, OM, SG, MPJ, and ALM carried out the histological determinations; Image extraction, analysis and correction were performed by ALM and SG; ALM, OM, MR, and VM carried out statistical analysis. ALM wrote the first draft of the manuscript; ALM, OM, MR, VM, PR, and RS contributed to the results and discussion. All authors read and approved the final manuscript.

## Conflict of interest

The authors declare no conflict of interest.

## Funding

This work is part of the project ’Identification des cibles Biochimiques et Histologiques impliquées dans la Réponse du maïs au déficit hydrIQUE (IBHERIQUE),’ funded by INRAE’s Department of Biology and Plant Breeding. Besides, Financial support has been provided by "PRIMA, a program supported by the European Union under H2020 framework program, and PCI2021-121912 funded by MCIN/AEI/ 10.13039/501100011033; and the Spanish Ministerio de Innovación y Universidades (MCIU), the Agencia Estatal de Investigación (AEI) and the European Fund for Regional Development (FEDER), UE (project code PID2019-108127RB-I00). ALM’s postdoctoral contract was granted by Xunta de Galicia’s program “Axudas de apoio a etapa de formación postdoutoral”.

## Data availability

Data supporting the findings of this study are available upon request from the corresponding author.

---

**Suplementary Figure 1:**
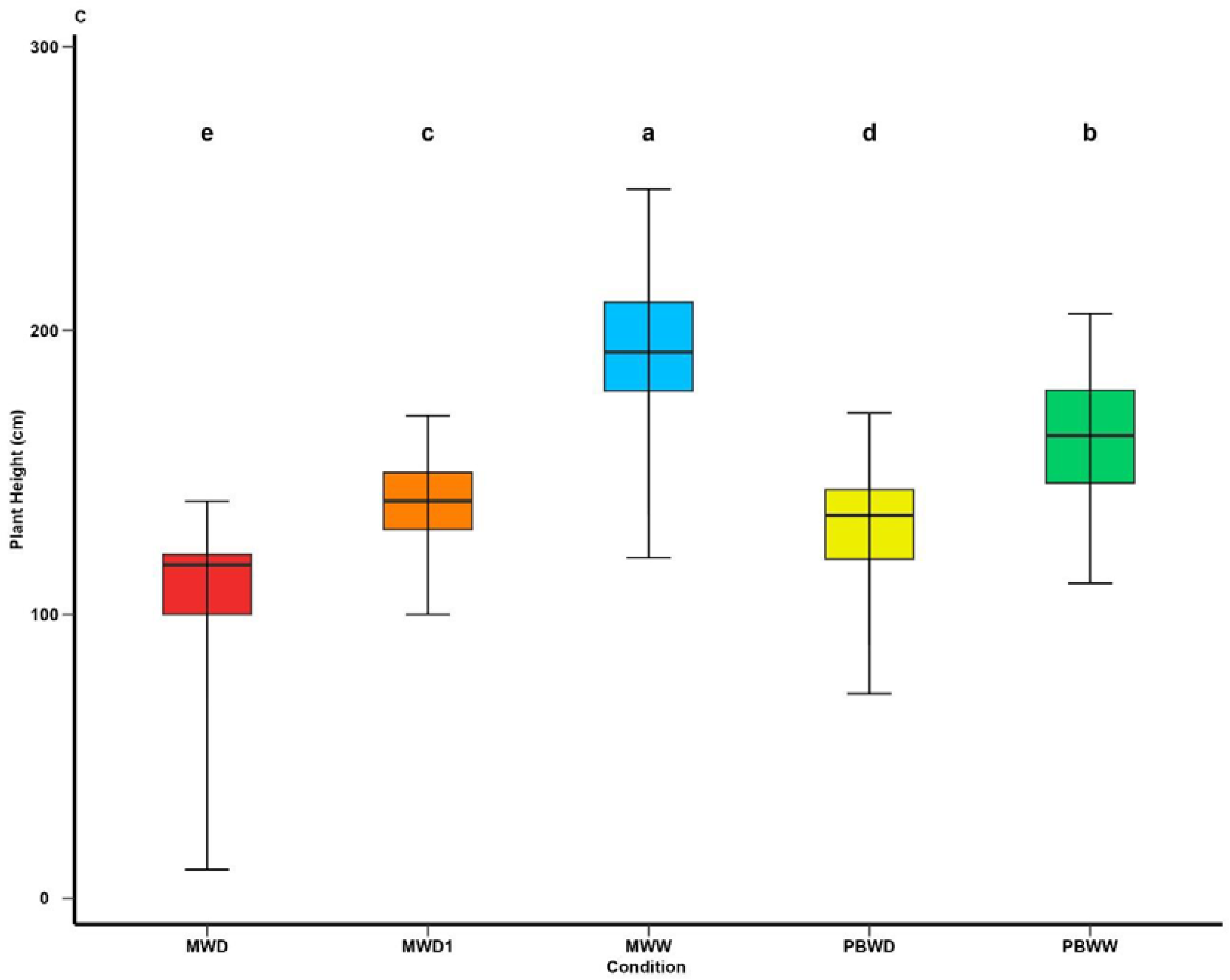
Means comparison of Plant Height under different irrigation treatments. Different letters indicate significant differences between conditions (p < 0.05).

## Notes

### Competing Interest Statement

The authors have declared no competing interest.

